# Deep learning-based approach for the characterization and quantification of histopathology in mouse models of colitis

**DOI:** 10.1101/2022.05.12.491690

**Authors:** Soma Kobayashi, Jason Shieh, Ainara Ruiz de Sabando, Julie Kim, Yang Liu, Sui Y. Zee, Prateek Prasanna, Agnieszka B. Bialkowska, Joel H. Saltz, Vincent W. Yang

## Abstract

Inflammatory bowel disease (IBD) is a chronic immune-mediated disease of the gastrointestinal tract. While therapies exist, response can be limited within the patient population. As such, researchers have studied mouse models of colitis to further understand its pathogenesis and identify new treatment targets. Although bench methods like flow cytometry and RNA-sequencing can characterize immune responses with single-cell resolution, whole murine colon specimens are processed at once. Given the simultaneous presence of colonic regions that are involved or uninvolved with abnormal histology, processing whole colons may lead to a loss of spatial context. Detecting these regions in hematoxylin and eosin (H&E)-stained colonic tissues offers the downstream potential of quantifying immune populations in areas with and without disease involvement by immunohistochemistry on serially sectioned slides. This could provide a complementary, spatially-aware approach to further characterize populations identified by other methods. However, detection of such regions requires expert interpretation by pathologists and is a tedious process that may be difficult to perform consistently across experiments. To this end, we have trained a deep learning model to detect ‘Involved’ and ‘Uninvolved’ regions from H&E-stained colonic slides across controls and three mouse models of colitis – the dextran sodium sulfate (DSS) chemical induction model, the recently established intestinal epithelium-specific, inducible *Klf5*^Δ*IND*^ (*Villin-CreER*^*T2*^*;Klf5*^*fl/fl*^) genetic model, and one that combines both induction methods. The trained classifier allows for extraction of ‘Involved’ colonic regions across mice to cluster and identify histological classes. Here, we show that quantification of ‘Involved’ and ‘Uninvolved’ image patch classes in swiss rolls of colonic specimens can be utilized to train a linear determinant analysis classifier to distinguish between mouse models. Such an approach has the potential for revealing histological links and improving synergy between various colitis mouse model studies to identify new therapeutic targets and pathophysiological mechanisms.

## Introduction

Inflammatory bowel disease (IBD) is a state of chronic intestinal inflammation and is comprised of two major subtypes, Crohn’s disease (CD) and ulcerative colitis (UC). IBD afflicts approximately 1.6 million Americans, and as many as 70,000 new cases are diagnosed each year (1). In addition to intestinal symptoms such as abdominal pain and diarrhea, patients’ quality of life can be severely affected by extraintestinal manifestations such as arthritis, ankylosing spondylitis, erythema nodosum, pyoderma gangrenosum, iritis, uveitis, and primary sclerosing cholangitis (2). Furthermore, CD has been linked to increased risk for cancer, pulmonary, gastrointestinal, genital, and urinary tract diseases (1), and UC is associated with an increased risk for colorectal cancer (3). Classically, CD and UC have different patterns of intestinal involvement. CD pathology is discontinuous and can affect any area along the gastrointestinal tract, while UC has a continuous distribution that is limited to the colon (4). Both diseases can have uninvolved intestinal regions. In CD, affected areas are termed ‘skip lesions’ as they are interspersed with uninvolved regions, and in UC, disease often does not involve the whole large intestine.

Mouse models of colitis have been heavily studied to understand disease pathogenesis and identify new treatment targets. One of the most used, in part due to the relatively simple method of induction and replicability, has been the dextran sodium sulfate (DSS) chemical induction model. DSS is provided to mice dissolved in drinking water for seven days to cause acute colitis (5). Groups have utilized methods such as flow cytometry and single-cell RNA sequencing to characterize the immune responses in these mice (6, 7). However, these methods require the processing of whole murine colons at once. As DSS-treated mice have been reported to have a predominantly distal injury pattern (8, 9), such an approach may not be sufficient to fully capture intracolonic heterogeneity. In addition to this injury pattern, Kolachala et al. reported different cytokine profiles between the proximal and distal colon (8). There is thus a likely benefit to resolving differences between this distal niche from the typically uninvolved proximal colon in DSS-treated mice. Additionally, our group has recently established the *Klf5*^Δ*IND*^ colitis mouse model (10, 11). We have also observed the simultaneous presence of colonic regions that are involved and uninvolved with disease in these mice. Therefore, we were motivated to develop an automated method to histologically identify these areas across these mouse models.

Convolutional neural networks (CNNs) have shown much promise in biomedical image classification tasks due to their ability to associate visual patterns with image labels (12-14). Provided that many histopathological labels are defined in part by visual patterns that pathologists have consistently seen for certain clinical diseases, CNNs are well-suited for such tasks. In practice, histological slides are digitized into gigapixel resolution whole slide images (WSIs). Due to the size of these WSIs, they are broken up into smaller, equally-sized image patches to which CNNs are applied. In this study, we sought to address whether CNNs could be incorporated in a pipeline to characterize intracolonic heterogeneity within mouse models. Although preclinical animal models are frequently utilized to identify novel protein and genomic targets, use of histology is often limited to confirming presence or lack of certain markers without a focus on the morphological and cellular patterns. As with the ‘villous’ or ‘adenomatous’ polyps, histological patterns are clinically significant and a manifestation of cellular and sub-cellular pathophysiological mechanisms. There is therefore a likely value to characterizing histological patterns in mouse models and tying their presence to molecular characterizations. Adaptation of histological phenotypes to routinely collected clinical specimens has the potential to transfer knowledge gained from animal models for more spatially-granular characterizations in a non-invasive manner.

A CNN pipeline to detect intracolonic heterogeneity could provide a histological readout to aid in the characterization of colitis mouse models and the evaluation of treatment efficacies. While this could be accomplished via pathologist’s interpretation, doing so for many samples is tedious and may be difficult to perform in a consistent manner across experiments and over time. Additionally, reliable identification of such regions is the first step towards correlating immunohistochemically identified cell populations to regions with and without abnormal histology. This could provide a spatial context-aware characterization of immune responses. Lastly, as we show in this study, a classifier trained for this task can extract diseased image regions across a cohort of mice. Involved region extraction allows for clustering approaches to identify histological classes, the variable presence of which can be utilized to distinguish between mouse models with an overall F1 score of 95.75%.

## Results

### Archived Mouse Cohort Background

To train our models, we utilized archived, formalin-fixed paraffin-embedded (FFPE) swiss-rolled colons (15). These were collected from three colitis mouse models and appropriate controls. The colitis model treatment schedules are detailed in S1A Fig. The first mouse model is the recently established *Klf5*^Δ*IND*^ (*Villin-CreER*^*T2*^*;Klf5*^*fl/fl*^) genetic model, wherein inducible intestinal epithelium-specific knockout of *Klf5* upon five days of intraperitoneal (IP) tamoxifen (TAM) injections disrupts epithelial barrier function and causes colitis (5T-*Klf5*^Δ*IND*^) (10, 11). For this study, we utilized female mice due to the higher rate of *Klf5* knockout relative to males (11). As TAM is dissolved in corn oil (CO), we also performed five days of IP CO injections in *Klf5*^Δ*IND*^ mice as a control (5C-*Klf5*^Δ*IND*^). Additionally, to provide controls for the genotypes and the TAM injections, we have also collected various combinations of controls with five days of IP CO and five days of IP TAM injections across *Klf5*^Δ*IND*^, *Klf5*^Δ*IND/+*^ (*Villin-CreER*^*T2*^*;Klf5*^*fl/+*^), and *Klf5*^*WT*^ mice (*Villin-CreER*^*T2*^*;Klf5*^*+/+*^) (S1 Table).

The second mouse model is the chemical dextran sodium sulfate (DSS) induction model. To control for the injections in our 5T-*Klf5*^Δ*IND*^ model, *Klf5*^Δ*IND/+*^ mice received five days of IP CO injections, which were then followed by seven days of 2.5% DSS in drinking water. We have seen that the histology mirrors that in non-injected *Klf5*^*WT*^ mice treated with DSS (S1B Fig). For the third model, we performed a combined induction. As homozygous 5T-*Klf5*^Δ*IND*^ mice exhibit increased mortality in an 18-day period after induction (11), we injected heterozygous *Klf5*^Δ*IND/+*^ mice with TAM for five days IP, then followed with seven days of 2.5% DSS in drinking water (5T-*Klf5*^Δ*IND/+*^*+* DSS). Although we have yet to fully characterize the combined induction model, we wanted to examine the histology in these mice and evaluate whether our computational pipeline could still distinguish the single-type induction models even in the presence of these mice in the cohort. In total, we have 48 mice in the archived mouse cohort, and sample numbers are available in S1 Table.

Of note, the colons extracted from these colitis models contain areas that are involved or uninvolved with disease. Manually selected regions from representative whole slide images (WSIs) are shown in Fig 1A. Swiss rolls allow for the visualization of a whole mouse colon on a single glass side. In these preparations, the center is the proximal end, and the colon tissue can be traced distally towards the outer portion of the swiss roll. Although not utilized in this study, we have also provided WSIs of additional controls and comparisons (5C-*Klf5*^Δ*IND/+*^ and 5T-*Klf5*^Δ*IND/+*^) for the combined induction model (S1C Fig). Clinical scoring metrics combining weight loss, stool consistency, and fecal blood (16) show significant increases for 5T-*Klf5*^Δ*IND/+*^ *+* DSS mice relative to 5C-*Klf5*^Δ*IND/+*^ + DSS and control mice (S1D Fig). Combined induction mice trend lower on histological scoring relative to 5C-*Klf5*^Δ*IND/+*^ + DSS mice (S1E Fig), mainly due to decreased extent of ulceration along the colon.

**Figure 1.**
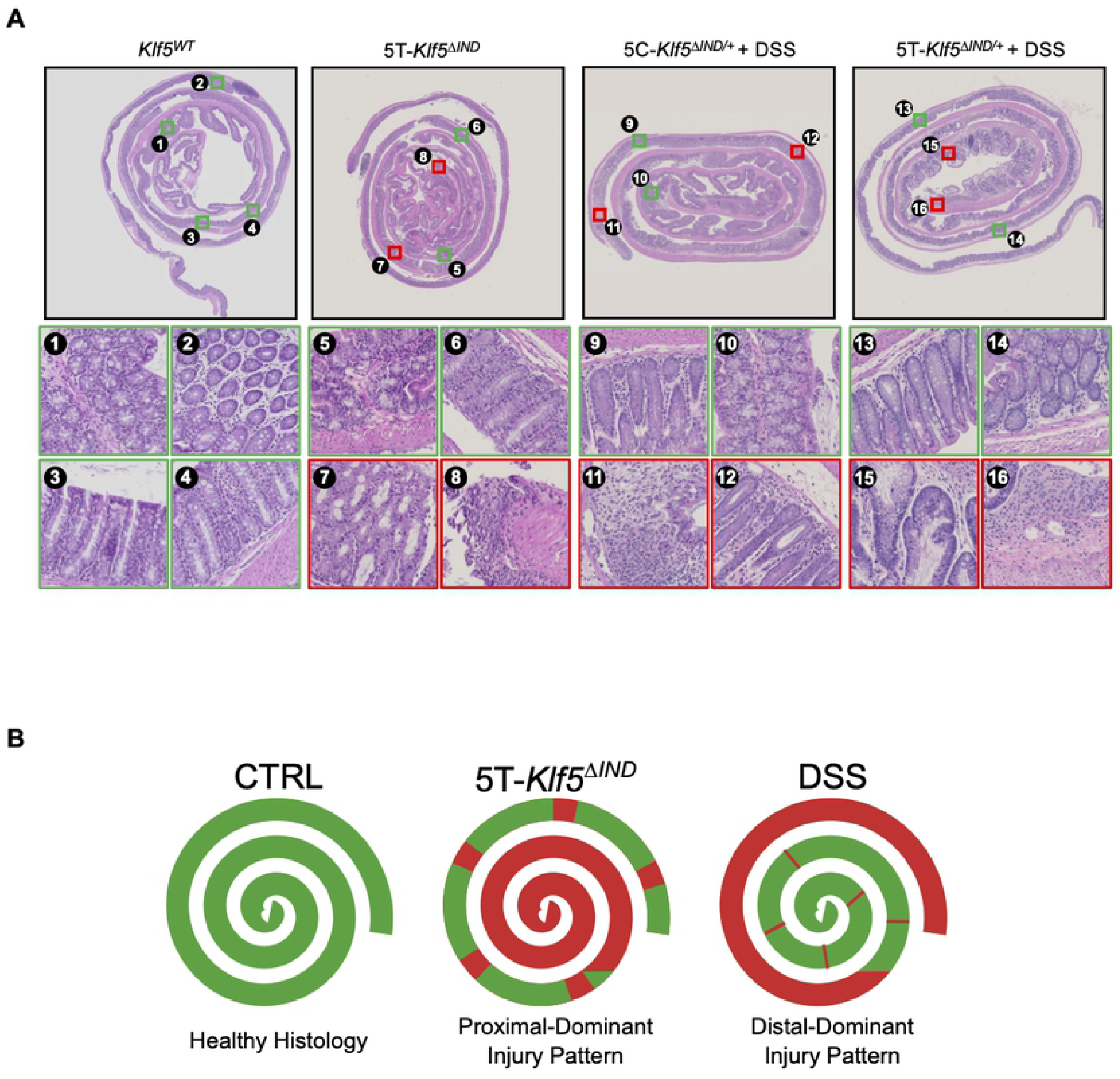
Colitis Mouse Models Exhibit Regional Heterogeneity. A) Representative whole slide images (WSIs) of swiss-rolled colons from *Klf5*^*WT*^ (*Villin-CreER*^*T2*^*;Klf5*^*+/+*^) control and colitis mouse models. Manually selected regions that are ‘Involved’ (red) and ‘Uninvolved’ (green) are shown for colitis mouse model samples. 8, 11, and 16 show areas with crypt dropout. 7 and 12 are examples of crypt dilation. 15 is an example of distorted glands. B) Reported patterns of injury in literature for colitis mouse models.

While the regional heterogeneity in our colitis models is visually apparent (Fig 1A), identification of these regions requires a pathologist’s inspection. This can be difficult at times to perform objectively and especially over many samples. Pathologist inter- and intra-observer variability has been reported in various contexts, such as lymph node counting (17), atypical ductal hyperplasia diagnosis (18), and follicular thyroid carcinoma diagnosis (19). An automated pipeline to identify murine colonic regions involved and uninvolved with disease would provide benefits in objective reproducibility and speed. Furthermore, reliable, automated identification of these areas may open the door for more spatially motivated, histological characterizations. A goal of this pipeline is to capture the reported proximal and distal injury patterns in the 5T-*Klf5*^Δ*IND*^ (11) and DSS models (8, 9), respectively (Fig 1B). Following detection, involved image regions across mice can be extracted to cluster and reveal pathologies specific to mouse models. To this end, we have trained an ‘Involved’ versus ‘Uninvolved’ region classification model that generates predictions over mucosal and submucosal regions of swiss-rolled murine colonic tissues.

### ‘Involved’ versus ‘Uninvolved’ Classifier Training

For our classifier, we used the ResNet-34 (RN-34) architecture (20). ResNet is considered a landmark architecture for the introduction of skip connections. Deep networks introduce a high number of non-linear mathematical operations through the addition of many layers, which can complicate the gradient descent operations that allow the neural network to “learn”. Gradient explosion and vanishing can occur when calculated gradients are too large or small and will impede learning (21). Skip connections allow gradient descent to skip layers of the network where this occurs, allowing for continued training. This has allowed for a deep, 34-layer ResNet that can extract even higher dimensional features from images, a powerful quality for H&E-patch classification. In this study, we trained RN-34 models that were pretrained on ImageNet, a large dataset with natural images comprising 1000 classes (22).

Our classifier was trained in a two-phase approach (S2A-B Fig). All WSIs across our archived mouse cohort were downsampled by a factor of 8 and tiled into equally sized 224×224 pixel patches. The initial phase trained a classifier with ground truth patch labels corresponding to mouse colitis status (S2A Fig). As such, all patches from colitis mice were labeled ‘Colitis’ and those from control mice were labeled ‘Control’. Since this labeling process does not account for the intracolonic heterogeneity observed in our colitis models (Fig 1A), we performed further feature extraction to extract the high dimensional patch representations learned during the first phase of training (S2B Fig). K-means clustering on these representations across all archived mouse cohort samples identified five patch classes (S2C-D Fig). Three of these classes (‘Crypts’, ‘Lightly Packed’, and ‘Rosettes’) were visually determined and validated with a pathologist to be uninvolved, while two of these classes (‘Mixed Pathology’, ‘Distorted Glands’) were involved with abnormal histology.

In addition to the visual evaluations, we calculated the proportion of each mouse’s patches that corresponded to each of these classes. Two of the three qualitatively uninvolved classes were significantly enriched in our control mice, while both of our involved classes were significantly enriched in colitis mice (S2D Fig). The lack of significance for the ‘Rosettes’ class is attributed to the higher proportion of patches of this class in DSS-treated mice relative to the other colitis models (S2E Fig). This is likely due to the predominantly distal injury pattern in DSS-treated mice (8, 9), as rosette structures are more prevalent in the proximal colon where abnormal histology is less common. Since mucosal folds are found in proximal murine colon (23, 24), the increased prevalence of rosette structures is due to sectioning at these regions. In the second phase of training (S2B Fig), these k-means classes were utilized as patch-level ground truth labels based on their qualitative descriptions and quantitative enrichment in colitis or control mice.

In both phases of training, 70% of total patches were used to train the model, 10% as a mid-training performance validation set, and 20% as a held-out test set. Our final model exhibited an overall F1 score of 90.1% when predictions were compared to our method-generated ground truth labels on the held-out test set (Fig 2A). This F1 score was an improvement from the overall F1 score of 77.6% for the model from the first phase of training with colitis status ground truth labels (S2F Fig). The first phase model exhibited a relatively lower F1 score of 68.87% when attempting to predict patches from colitis mice. This performance is likely explained by the presence of uninvolved patches extracted from our colitis mice labeled as ‘Colitis’ and motivated our second phase of model training. Additionally, when applied to 171 patches with ‘Involved’ and ‘Uninvolved’ ground truth labels provided by a pathologist, our classifier exhibited an overall F1 score of 81.41% and again showed improvement from the model from the first phase of training (S2G Fig).

**Figure 2.**
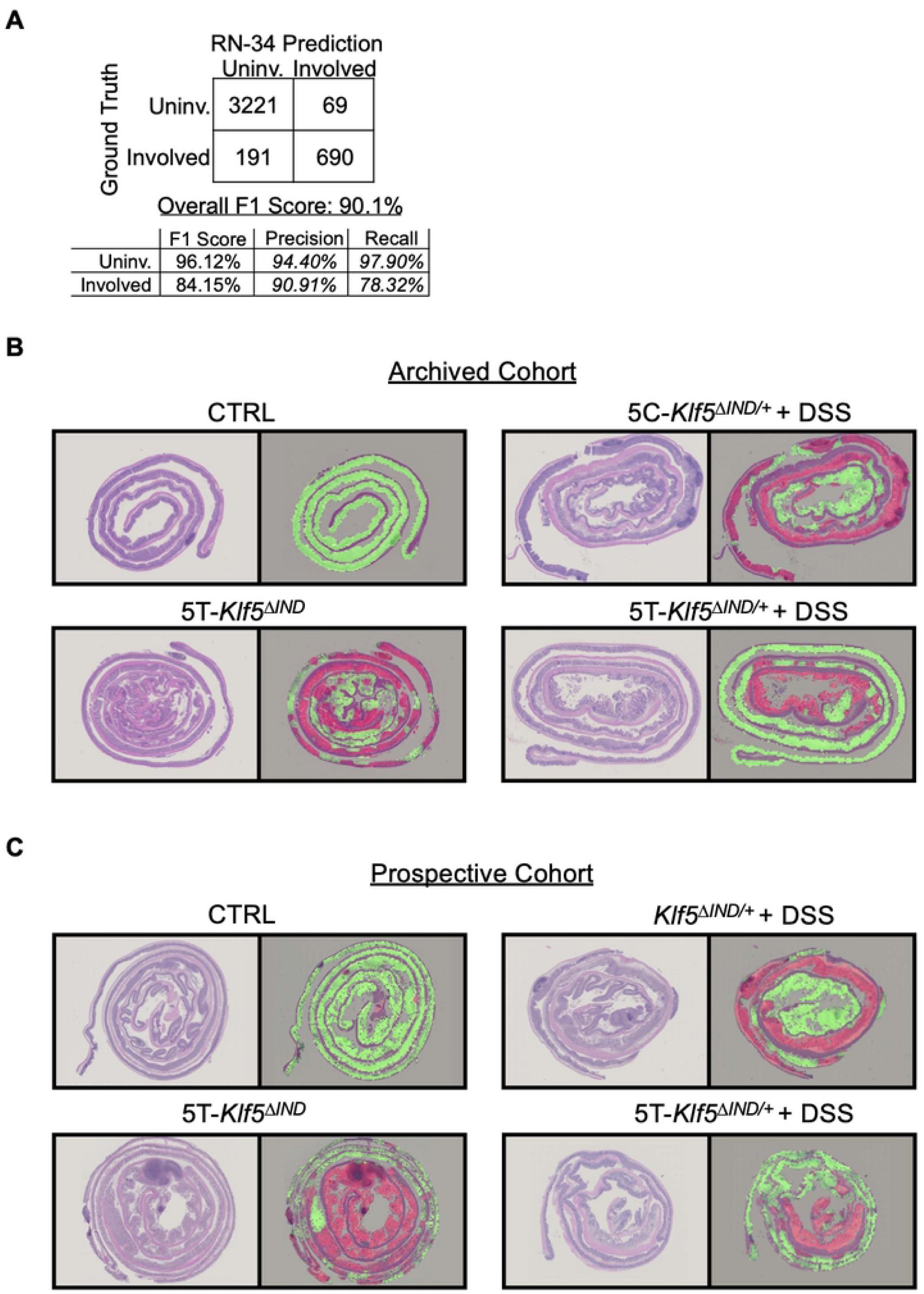
H&E Model Performs ‘Involved’ versus ‘Uninvolved’ Classifications on Swiss-Rolled Colons. A) Confusion matrix showing final model predictions on patches from test set mice. From the 48 mice in our archived cohort (S1 Table), 34 mice (14281 patches, ∼70% of total) were used for model training, 5 mice were used for mid-training validation (2192 patches, ∼10% of total), and 9 mice (4171 patches, ∼19%) were used as a final independent test set. B) Original input H&E-stained swiss rolls are shown with corresponding ‘Involved’ (red) versus ‘Uninvolved’ (green) classifier overlays for test set mice from archived mouse cohort. C) Original input H&E-stained swiss rolls and corresponding overlays for mice from prospectively treated mouse cohort. For both B) and C), red and green overlay colors correspond to regions with at least 50% model prediction confidence.

### Small Patch Classifier Training for Preprocessing

We initially used simple thresholding of pixel intensities across each patch’s red, green, and blue color channels to filter out patches with too much background and muscle. However, we observed that this approach was not robust and trained an additional RN-34 classifier pretrained on ImageNet on smaller 32×32 pixel patches to classify between ‘Background’, ‘Muscle’, ‘Tissue’, and ‘Submucosa’ classes (S3 Fig). From 8 mice (2 each of control and of the three mouse models), we extracted 100 patches for each of the ‘Background’, ‘Muscle’, ‘Tissue’, and ‘Submucosa’ classes (S3A Fig). Our model was trained with these ground truth labels and applied to a test set of patches from 4 different mice as training (1 each of control and of the 3 mouse models, 100 patches for each class). This model achieved an overall F1 score of 93.5% (S3B Fig). We also generated qualitative overlays of our small patch classifier’s prediction on test set mice (S3C Fig). Additionally, we opted to implement this small patch classifier within our patch filtering process. All patches with >65% of area corresponding to regions predicted to be ‘Background’ or ‘Muscle’ are filtered out during the patch extraction process (S4 Fig).

### ‘Involved’ versus ‘Uninvolved’ Overlay Generation

To provide visual outputs for classifications, we generated overlays for our held-out test set mice (Fig 2B). Areas with at least 50% confidence of ‘Involved’ and ‘Uninvolved’ predictions are overlayed onto the input H&E WSIs. Overlapping patches were extracted from all downsized WSIs. This was done by taking initially extracted patch locations and iterating 20 pixels in each direction (up/down/left/right). Overlapping patch extraction led to an increase of ∼450 to ∼200,000 patches per mouse after repeating filtering to eliminate patches with too much muscle or background. Our classifier was applied to each overlapping patch, and ‘Involved’ and ‘Uninvolved’ prediction confidences were averaged at every pixel in the WSI. All regions predicted by our Small Patch Classifier to be ‘Background’ and ‘Muscle’ were eliminated from final overlays.

### Prospective Mouse Cohort

To test the robustness of our ‘Involved’ versus ‘Uninvolved’ classifier, we treated and collected H&E-stained, FFPE swiss-rolled colons from additional mice. Specifically, this cohort allowed us to assess whether our approaches would work regardless of who treated mice, collected swiss rolls, and performed H&E staining. To complement the 48 mice in our archived mouse cohort, we treated 24 additional mice. This consisted of eight controls (no injections, put on normal drinking water), eight *Klf5*^Δ*IND/+*^ DSS-treated mice with no injections, five 5T-*Klf5*^Δ*IND*^, and three 5T-*Klf5*^Δ*IND/+*^ *+* DSS mice (S1 Table). We elected to utilize non-injected DSS-treated mice to confirm that our model properly captures the histology in these mice even without five days of corn oil injections preceding the DSS. We applied our approach to these prospective samples and successfully generated ‘Involved’ versus ‘Uninvolved’ overlays (Fig 2C).

### Patch Class Discovery

To build upon the binary classifications generated by our final model, we aimed to define more specific image patch classes. Analogous to our second phase of model training (S2B Fig), we utilized our final ‘Involved’ versus ‘Uninvolved’ classifier to extract high dimensional representations for patches in our dataset. Specifically, two datasets (one with all ‘Involved’ patch representations and one with all ‘Uninvolved’ patch representations) were collected. Given the high number of patches, principal component analysis (PCA)-based dimensionality reduction was then performed on each set of patch representations before k-means clustering. Clustering on ‘Uninvolved’ patches from our archived mouse cohort identified three classes: ‘Crypts’ – test-tube-like perspective of crypt structures, ‘Lightly Packed’ – more space between crypts often accompanied by immune cell nuclei, and ‘Rosettes’ – Ring-like cross-section perspective of crypts (Fig 3A-B). Of note, the discovered ‘Uninvolved’ patch classes matched those utilized as ground truth labeling during our second phase of classifier training (Fig S2C).

**Figure 3.**
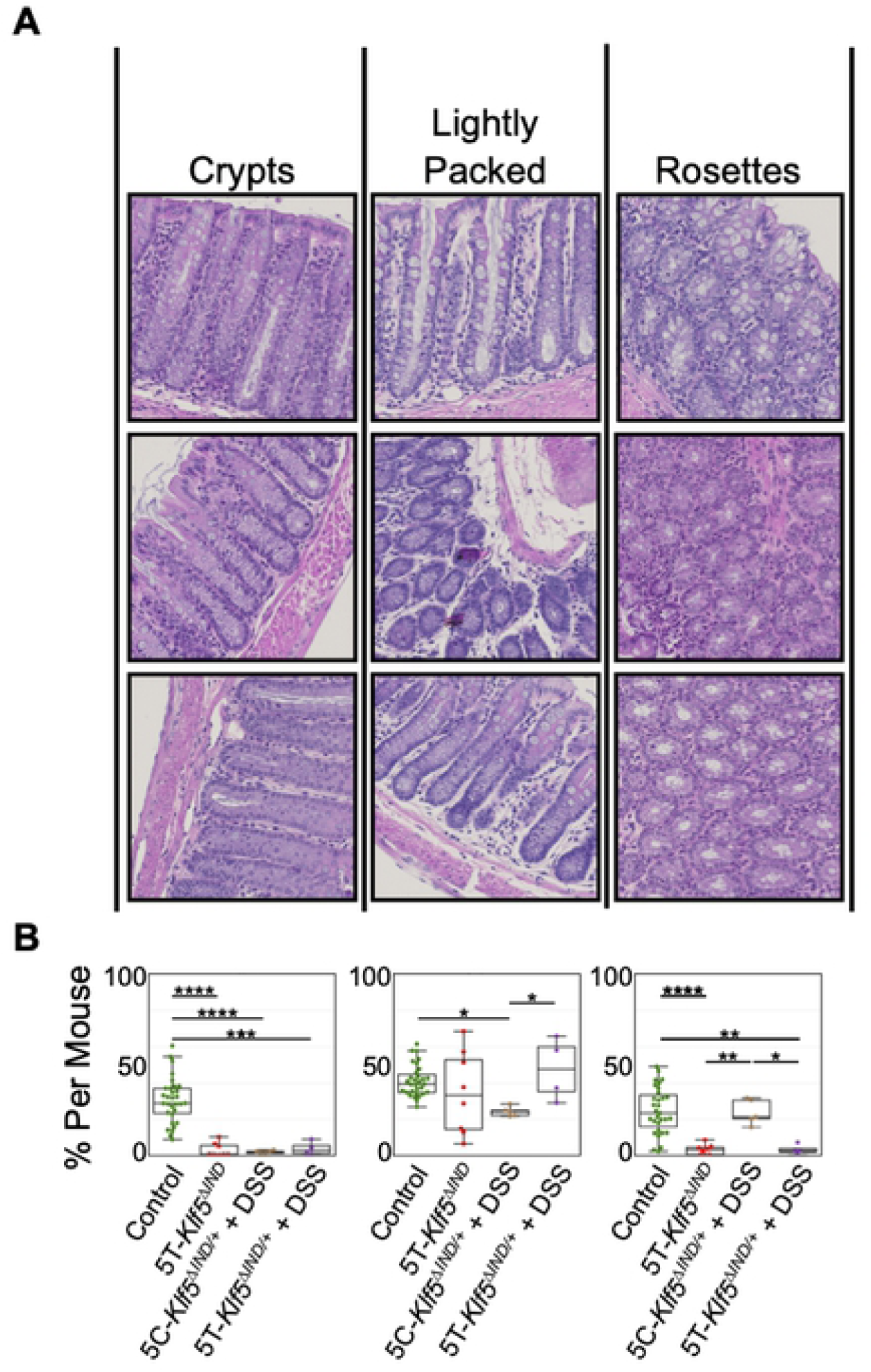
Discovered ‘Uninvolved’ Patch Classes are Enriched in Control Mice and Show Shifts in Prevalence Across Colitis Mouse Models. A) ‘Uninvolved’ patch classes with qualitative class labels. Patches most representative of class labels are shown. B) One-way ANOVA shows that these ‘Uninvolved’ patch classes are found in differing portions across colitis mouse models and controls. *p<0.05, **p<0.01, ***p<0.001, ****p<0.0001.

While the ‘Crypts’ class was significantly enriched in control mice relative to all the colitis mouse models, this was not true for the other two classes. Specifically, the ‘Rosettes’ class was significantly enriched in control mice relative to 5T-*Klf5*^Δ*IND*^ and 5T-*Klf5*^Δ*IND/+*^ + DSS mice, but not in 5C-*Klf5*^Δ*IND/+*^ + DSS mice (Fig 3B). In addition, the ‘Rosettes’ class was significantly enriched in the 5C-*Klf5*^Δ*IND/+*^ + DSS mice relative to the other two colitis models. This is again likely due to the predominantly distal injury pattern in DSS-treated mice (8, 9). This injury pattern is also depicted in our ‘Involved’ versus ‘Uninvolved’ overlays (Fig 2B-C). Our DSS-treated mice show more ‘Uninvolved’ regions in the rosettes-enriched proximal colon (center of swiss rolls) where disease is less common. In the 5T-*Klf5*^Δ*IND/+*^ + DSS mice, however, DSS-treatment no longer causes a distal predominant injury pattern (Fig 2B-C). While the mechanism of DSS-induced colitis has not been completely elucidated, the effects are believed to be dependent on tissue penetration of the chemical leading to disruption of the intestinal epithelial monolayer and barrier integrity (25). DSS has a variable molecular weight from 5 to 1400 kDa, and administration of forms of 500 kDa and higher do not induce colitis (26). As such, the shift in the distal injury pattern of 5C-*Klf5*^Δ*IND/+*^ + DSS mice to the proximal one in 5T-*Klf5*^Δ*IND/+*^ + DSS mice may be a function of increased proximal tissue permeability from TAM injection-induced heterozygous knockout of KLF5 that promotes greater tissue penetrance of DSS.

The ‘Lightly Packed’ class was significantly enriched in control mice relative to 5C-*Klf5*^Δ*IND/+*^ + DSS mice but not the other two colitis mouse models. This likely relates to the increased prevalence of abnormal proximal histology in the 5T-*Klf5*^Δ*IND*^ and 5T-*Klf5*^Δ*IND/+*^ + DSS models relative to 5C-*Klf5*^Δ*IND/+*^ + DSS mice (11) (Fig 2B-C). However, the ‘Crypts’, which are found more distally, significantly decreased in all colitis mouse models relative to controls. Thus, an additional explanation is that the ‘Lightly Packed’ regions in these mice sit on the decision border, representing an accumulation of immune cells without sufficient levels of abnormal histology to garner an ‘Involved’ prediction. While our 5T-*Klf5*^Δ*IND/+*^ + DSS model exhibited a significantly higher clinical score than our 5C-*Klf5*^Δ*IND/+*^ + DSS mice (S1D Fig), they trended lower in histological scoring (S1E Fig). This may reflect the limitation of histological scoring schemes that rely upon simple detection and counting of pre-defined pathological findings without accounting for lower grade abnormalities in other regions. Thus, in our 5T-*Klf5*^Δ*IND*^ and 5T-*Klf5*^Δ*IND/+*^ + DSS mice, the presence of ‘Lightly Packed’ patches may indicate an immune response that is present but either protective or insufficient to generate overt abnormalities. A future direction is thus to leverage this property of our pipeline to consider these borderline regions in the grading of swiss rolls.

The ‘Mixed Pathology’ k-means class identified during our second phase of model training (S2B-D Fig) motivated us to apply this approach to ‘Involved’ patches from across the archived mouse cohort to identify specific types of abnormal histology. K-means clustering on these ‘Involved’ patches identified four pathologic patch classes (Fig 4A-B). By visual inspection and validation by a pathologist, these were classified as ‘Inflammatory’ (Milder, more heterogeneous histological phenotype with an influx of immune cell nuclei), ‘Crypt Dropout’ (loss of epithelium and displacement by stroma and immune cells), ‘Crypt Dilation’ (expansion of crypt lumen spaces), and ‘Distorted Glands’ (distortion of crypt structures). Of note, the ‘Distorted Glands’ class persisted from the ground truth classes used in our second round of model training (S2C-D Fig), but the ‘Mixed Pathologies’ class was replaced by the other classes. Notably, immune cell infiltration, crypt dilation, and gland distortion are all histological features that have been observed in human IBD (27-29). Furthermore, these ‘Involved’ patch classes were found to be present in significantly different proportions across our mouse models and controls (Fig 4B). Specifically, 5C-*Klf5*^Δ*IND/+*^ + DSS mice were enriched in the ‘Inflammatory’ and ‘Crypt Dropout’ classes, 5T-*Klf5*^Δ*IND*^ mice in the ‘Inflammatory’ and ‘Crypt Dilation’ classes, and 5T-*Klf5*^Δ*IND/+*^ + DSS mice in the ‘Crypt Dropout’ and ‘Distorted Glands’ classes (Fig 4B).

**Figure 4.**
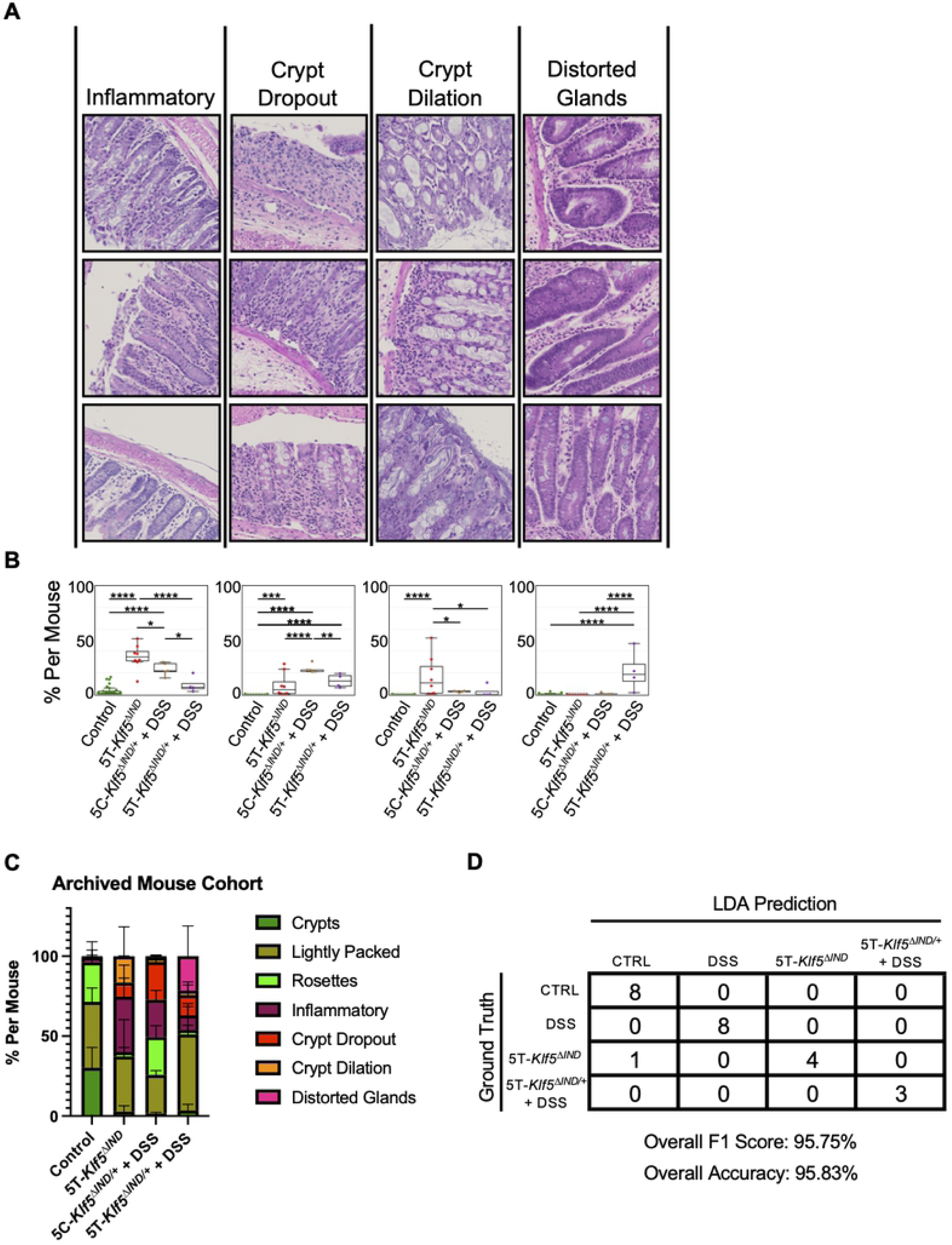
Discovered ‘Involved’ K-Means Patch Classes Can Be Utilized by Machine Learning Classifier to Predict Mouse Model. A) ‘Involved’ patch classes with qualitative class labels. Patches most representative of class labels are shown. B) One-way ANOVA shows that these ‘Involved’ patch classes are found in differing portions across colitis mouse models and controls. *p<0.05, **p<0.01, ***p<0.001, ****p<0.0001. C) Stacked bar plot of total ‘Uninvolved’ and ‘Involved’ k-means patch class proportions across archived mouse cohort. Error bars show mean with standard deviation. D) Confusion matrix for linear determinant analysis (LDA) classifier mouse model prediction on prospective mouse cohort. LDA model was trained on the original 48 mice cohort of archived tissues.

Visualizing patch class proportions by stacked bar plots helps capture and summarize these findings (Fig 4C). While all three ‘Uninvolved’ patch classes are present in control mice, ‘Crypts’ patches fall significantly for all three colitis mouse models, while ‘Lightly Packed’ patches persist. However, the presence of ‘Rosettes’ are maintained in our 5C-*Klf5*^Δ*IND/+*^ + DSS mice. The ‘Inflammatory’ and ‘Crypt Dilation’ patch classes are most prevalent in our 5T-*Klf5*^Δ*IND*^ mice. Given the shorter treatment course of this colitis mouse model (5 days), these patch classes may indicate a more acute finding. On the other hand, the ‘Crypt Dropout’ class may be less acute as prevalence is higher in our longer treatment course 5C-*Klf5*^Δ*IND/+*^ + DSS (7 days) and 5T-*Klf5*^Δ*IND/ +*^ + DSS mice (12 days). Similarly, our 5T-*Klf5*^Δ*IND/+*^ + DSS mice, which have 12 days of total colitis induction (five days of TAM injections, seven days of DSS, S1 Table), are the only model with appreciable presence of the distorted glands class. Distorted glands are one of the histological markers of chronic inflammation in human IBD (30) and exemplifies how this type of pipeline can histologically categorize colitis mouse models by which facets of human disease they recapitulate. In addition, partnering this approach with immunohistochemical staining can shed further light on the relative contributions of length of colitis induction and immune cell presence in causing various types of histological abnormalities.

### Prediction of Mouse Model from H&E Inputs

Finally, we sought to address whether the variable presence of ‘Uninvolved’ patches and ‘Involved’ patch classes could be utilized to predict mouse model in our prospective mouse cohort. We thus formed the pipeline detailed in S5A Fig to utilize ‘Uninvolved’ patch and ‘Involved’ k-means patch class proportions to train a linear determinant analysis (LDA) classifier to predict mouse model.

The LDA classifier was trained on the archived mouse cohort. When applied to the prospective mouse cohort via the inference pipeline in S5B Fig, this LDA classifier predicted which mouse model each swiss roll came from with an overall F1 score of 95.75% (Fig 4D). Of note, the stacked bar plots of k-means patch classes in the prospective cohort (S5C Fig) mirrored what we observed in the archived mouse cohort (Fig 4C). As one of the 5T-*Klf5*^Δ*IND*^ mice was classified as control, we examined the input H&E and found healthier appearing histology relative to properly classified 5T-*Klf5*^Δ*IND*^ mice. This was apparent even from low magnification with decreased luminal space areas and more tightly packed crypts (S6 Fig). Although some ‘Involved’ areas were present in the 5T-*Klf5*^Δ*IND*^ mouse classified as control, crypts were more intact in these regions and accompanied with goblet cell presence. Goblet cells secrete mucus to provide a protective lining in the intestine, and decreased goblet cells are observed in CD and UC (31). As such, this pipeline can histologically identify outlier mice with weaker colitis induction and phenotypes.

## Discussion

This study sought to address the simultaneous presence of colonic regions that are histologically involved and uninvolved with abnormal histology within colitis mouse models. While these regions can be identified via careful inspection by a pathologist, doing so objectively over many samples is difficult and highly time-consuming. Deep learning models learn to associate class labels with implicit patterns in data and are well-suited for this task. A computational method to quantify the presence of abnormal histology can also serve as a histological readout for experiments performed on mice. One example would be to confirm the effect of treatments histologically as an accompaniment to clinical metrics like body weight, fecal blood, or stool consistency.

We have thus presented an ‘Involved’ versus ‘Uninvolved’ classifier for colons of DSS-treated, 5T-*Klf5*^Δ*IND*^, 5T-*Klf5*^Δ*IND/+*^ *+* DSS, and control mice. Specifically, we have utilized a two-phase training approach. The first phase uses mouse colitis status as patch ground truth labels, while the second phase incorporates feature extraction and clustering to generate improved, patch-level ground truth labeling. Importantly, our final model shows improved agreement with a pathologist relative to the model trained in just the first phase of the approach.

In a related study, Bédard et al. published a proof of concept application of convolutional neural networks for the microscopic scoring of acute inflammation in DSS colitis (32). They utilized a commercial artificial intelligence platform to detect muscle, normal mucosa, and acutely inflamed mucosa in H&E-stained murine colon WSIs. The authors calculated the ratio of acutely inflamed to total mucosa to assess the level of disease. Identifying these regions is an important step towards the microscopic grading of colitis samples. We have similarly trained our Small Patch Classifier to detect ‘Background’, ‘Muscle’, ‘Mucosa’, and ‘Submucosa’ classes from 32×32 pixel patches. In our work, the Small Patch Classifier was implemented to improve our patch filtering process and to focus our ‘Involved’ versus ‘Uninvolved’ classifier overlays onto the ‘Mucosa’ and ‘Submucosa’ regions. A future direction is to explore whether the Small Patch Classifier outputs can improve our ‘Involved’ versus ‘Uninvolved’ model performance.

In an alternate approach, Rogers et al. attempted to segment colitis lesions in H&E-stained WSIs across the DSS, CD45RBHi, and IL-10^-/-^ mouse models to grade disease severity (33). The authors experienced segmentation challenges due to variabilities in crypt morphologies and opted to generate a workflow based on CD3 immunohistochemical staining instead. Morphological variabilities likely increase more with disease relative to healthy histology. Here, we chose to train a patch-based ‘Involved’ versus ‘Uninvolved’ classifier. Including the task to classify ‘Uninvolved’ regions that should be more homogenous in morphology may help to separate out the ‘Involved’ regions. An additional benefit is that this has allowed us to perform clustering on just the ‘Involved’ patches to identify specific types of abnormal histology. Clustering on ‘Uninvolved’ patches also generated classes of normal histology. As with Rogers et al., we have also incorporated multiple mouse models of colitis in our study. This has allowed for comparisons showing quantitative enrichment of pathologic classes in the different mouse models.

Specifically, we applied feature extraction, PCA-based dimensionality reduction and k-means clustering on ‘Uninvolved’ patches to define ‘Crypt’, ‘Rosette’, and ‘Lightly Packed’ classes. Patches of the ‘Crypt’ class were significantly decreased in all colitis mouse models. The ‘Rosette’ and ‘Lightly Packed’ classes were present in variable proportions across mouse models, mirroring shifts in proximal and distal predominant patterning of injury. As the ‘Lightly Packed’ classes in colitis mice may represent regions of immune infiltration without overt abnormal histology, we plan to assess the capacity of our model to incorporate these regions in the grading of colonic histology.

From ‘Involved’ patches, we identified four classes of abnormal histology – ‘Inflammatory, ‘Crypt Dropout’, ‘Crypt Dilation’, and ‘Distorted Glands’. DSS-treated mice were enriched in the ‘Inflammatory’ and ‘Crypt Dropout’ classes, 5T-*Klf5*^Δ*IND*^ mice in the ‘Inflammatory’ and ‘Crypt Dilation’ classes, and 5T-*Klf5*^Δ*IND/+*^ + DSS in the ‘Crypt Dropout’ and ‘Distorted Glands’ classes. The proportion of patches per swiss roll classified as ‘Uninvolved’ and the four ‘Involved’ classes for each mouse was sufficient to train an LDA classifier to predict mouse model. Increased prevalence of the ‘Inflammatory’ and ‘Crypt Dilation’ classes in the shorter time course 5T-*Klf5*^Δ*IND*^ (5 days) and DSS-treated (7 days) mice may indicate these are more acute findings. Increases of ‘Distorted Glands’ in the combined induction model (12 days) and of ‘Crypt Dropout’ in the DSS-treated mice and combined induction model may reflect the more chronic nature of these histological classes. To the best of our knowledge, this is the first study to explore quantitation of pathologies across mouse models to invite further assessment of histology beyond qualitative descriptions. Additionally, this may help validate the capacity of mouse models to capture specific aspects of human disease. Detection of specific histologic abnormalities will likely provide finer grained characterizations for improved mappings to human IBD heterogeneity. To allow others to utilize the code, we have made available our prospective mouse cohort images and the full inference pipeline from WSI scaling and patch extraction to LDA inference.

Our pipeline thus essentially examines swiss rolls, detects and categorizes histology, then predicts mouse model. As the 5T-*Klf5*^Δ*IND*^ mice have a Th17-mediated immune response (11), while the DSS model has been characterized as Th1-, Th17-, and innate immunity-mediated (6, 34, 35), we have confirmed that the variable immune responses in these models are accompanied by differences in histological pathologies. The next step is to delineate the respective contributions of various immune populations to presence of specific pathologies.

Accordingly, we believe this pipeline is the start for a spatially motivated characterization of immune populations by histologic localization to ‘Involved’ and ‘Uninvolved’ regions. While methods like flow cytometry and single-cell sequencing can characterize immune responses with single-cell resolution (6, 7), whole tissues must be processed at once and spatial context is sacrificed. As a complement to these methods, a future direction is to map immune populations stained on serially sectioned slides to this regional heterogeneity as another approach to assessing immune cell functionality and subtypes. Residence of immune populations in areas with and without disease may provide another angle towards characterization. Since many studies assessing functionality rely on knockouts or expensive neutralizing antibody experiments (11, 36-40), such a pipeline may offer a cheaper, quicker alternative.

One limitation of this method is that current application is restricted to the mouse models included in this study. However, we believe this is an important first step to establish and promote the potential of computer vision methodologies at the bench, and specifically, within the context of colitis mouse models. Another future direction is therefore to amplify the capacity of this method by incorporating other colitis models within training, such as the 2,4,6-Trinitrobenzenesulfonic acid (TNBS), IL-10 knockout, adoptive cell transfer, and oxazolone models (41). Doing so would allow for the application of our method as a histological readout over a wider range of colitis mouse models.

More importantly, ‘Involved’ regions could then be extracted over an even more heterogeneous collection of colitis phenotypes. As these colitis models across literature differ in induction and flavor of immune responses, such an approach would open the door for synergy of knowledge gained from different mouse models. One possibility is that immune response characterizations, molecular approaches, and mechanistic studies focusing on pathophysiology could be linked to similar and dissimilar presence of types of histological findings across different murine colitis phenotypes. Notably, protein and genomic targets are often identified in preclinical animal models then evaluated for clinical value in patient specimens. Histological patterns identified in these animal models are driven by these same protein and genomic targets. Consequently, a future goal is to evaluate the clinical value and transferability of such histological phenotypes identified in animal models. This approach, therefore, has the potential to one day promote the discovery of novel therapeutic targets and pathways in IBD by serving as link between studies. As such, we believe that the integration of computational and computer vision approaches offers significant and exciting potential in bringing together the depth of human knowledge that has been gained from across colitis mouse models and in encouraging further collaboration across groups.

## Materials and Methods

### Mice

All studies involving mice were approved by the Stony Brook University Institutional Animal Care and Use Committee (IACUC; protocol #354918 and 245094). All mice carried an inducible Cre recombinase gene under the *Villin* promoter. These mice carried either two wild-type alleles of *Klf5* (*Villin-CreER*^*T2*^*;Klf5*^*+/+*^) or were heterozygous or homozygous for an additional *Klf5* allele flanked by loxP sites (heterozygous: *Villin-CreER*^*T2*^*;Klf5*^Δ*IND/+*^, homozygous: *Villin-CreER*^*T2*^; *Klf5*^Δ*IND/ΔIND*^).

Treatment schedules are detailed in S1A Fig and all treatment conditions are available in S1 Table. For all experiments, mice were eight to ten weeks old and female, as we have seen increased Cre recombination efficiency in these mice relative to males (11). Intraperitoneal (IP) injection of tamoxifen at 1mg/day dissolved in corn oil for 5 days was performed as previously described (11). Control mice received 5 days of IP corn oil injections. For dextran sodium sulfate (DSS) treatment, mice in the archived cohort received 2.5% DSS in drinking water. In the prospective cohort, DSS-treated mice received 3% DSS, as we observed this was the optimal concentration for experiments at the Stony Brook facilities. Combined induction mice received 2.5% DSS following 5 days of IP injection of tamoxifen at 1mg/day dissolved in corn oil.

### Whole Slide Image (WSI) Generation

FFPE swiss-rolled colons were sectioned onto glass slides and stained by hematoxylin and eosin (H&E) (15). These glass slides were then scanned and digitized at 40X magnification (0.17 µM/pixel) to a .vsi format by the Olympus VS120 Digital Virtual Slide System (VS120-L100-W). The files were converted to a tiff format for further downstream use.

### H&E ‘Involved’ versus ‘Uninvolved’ Classifier Training

For this study, we trained the ResNet-34 (RN-34) neural network, which is considered a landmark architecture for its introduction of skip connections (20). Briefly, skip connections allow the model to ‘skip’ parts of its architecture where training would typically be impeded due to gradient explosion or vanishing. We trained a RN-34 model that was pretrained on ImageNet, a large dataset with natural images comprising 1000 classes (22).

An overview schematic of our two-phase training approach is available in S2A-B Fig. The initial step is to downsize our WSIs by a factor of 8 and extract 224×224 pixel patches. Patches containing too little tissue are filtered out. The first phase, outlined in S2A Fig, utilizes mouse-level colitis induction status as ground truth to label extracted patches. As such, while our colitis mice show colonic presence of regions that are involved and uninvolved with disease (Fig 1A), this is not addressed here. Instead, all patches from mice with colitis induction are provided a ‘Colitis’ label, while all patches from control mice are provided a ‘Control’ label. Once labeled, 70% of patches were used to train the model, 10% of patches were used assess to performance mid-training, and 20% of patches were used as a held-out test set.

With the goal of generating more granular, patch-level ground truth labeling, we pursued the approach detailed in S2B Fig. We used our “Before k-means” as a feature extractor to convert all image patches in the archived mouse cohort dataset to high-dimensional 512-length numerical representations learned during the first phase of training in S2A Fig. K-means clustering on these representations generated patch classes (S2C Fig).

We then utilized these k-means clusters to generate new patch-level ground truth labels. As our control mice did not receive any colitis induction, we provided ‘Uninvolved’ ground truth labels to all patches from these mice. For our mice with colitis induction, we referred to the k-means clustering results. With this new ground truth labeling, we trained another RN-34 model pretrained on ImageNet using a 70/10/20% training/validation/test set split.

### Small Patch Classifier Training for Preprocessing

From our archived mouse cohort dataset WSIs, we extracted 32×32 pixel patches and trained an additional RN-34 classifier pretrained on ImageNet (S3 Fig). Our Small Patch Classifier predicts these patches as one of the ‘Background’, ‘Muscle’, ‘Tissue’, and ‘Submucosa’ classes.

### Tissue Map Generation and Patch Filtering

For every H&E-stained, swiss-rolled colon WSI, we first extract 32×32 pixel patches. The Small Patch Classifier is applied to each to generate a tissue map for every input H&E WSI (S4A Fig). The tissue map is of the same dimension as the input H&E where white areas represent ‘Tissue’ and ‘Submucosa’ classifications, while the black areas represent ‘Muscle’ and ‘Background’. For each subsequently extracted 224×224 pixel H&E patch, the corresponding area is extracted from the accompanying tissue map. Each patch is then thresholded to evaluate whether enough tissue or submucosa is present. Practically, we implement a 65% threshold. Patches with more than 65% of ‘Background’ or ‘Muscle’ regions are filtered out, while those with less are kept (S4B Fig).

### Overlap Patch Extraction and Overlay Generation

To reduce classification dependence for what patch happened to be extracted from a region in a WSI, we extracted overlapping patches. Beginning with existing patch locations, we iterated 20 pixels 10 times in each direction (up/down/left/right) and extracted new patches. We then again filtered out patches containing too little tissue. This led to an increase of ∼450 to ∼200,000 patches per mouse.

To generate overlays, our ‘Involved’ versus ‘Uninvolved’ classifier was applied to all patches, including overlapping patches. At every pixel in the original H&E, ‘Involved’ and ‘Uninvolved’ prediction confidences were averaged. As such, a greater degree of spatial context is incorporated for final classifications at each pixel. Red areas correspond to pixels with >50% confidence of ‘Involved’ predictions, while green areas correspond to pixels with >50% confidence of ‘Uninvolved’ predictions. Furthermore, all regions in tissue maps corresponding to ‘Muscle’ and ‘Background’ were eliminated from the overlay. This generated final outputs with overlays only over mucosal and submucosal areas (Fig 2B-C).

### Discovery of Patch Classes

Overview schematic is shown in S5A Fig. From across our archived mouse cohort, we extracted all patches, including overlapping patches, classified as ‘Involved’. We utilized our final ‘Involved’ versus ‘Uninvolved’ classifier as a feature extractor then performed PCA-based dimensionality reduction to 250 principal components (PCs) that captured 95% of variability. We then performed k-means clustering to identify pathologic patch classes. This process was also repeated for all patches classified as ‘Uninvolved’ in our archived mouse cohort dataset to identify non-pathologic patch classes. PCA-based dimensionality reduction for ‘Uninvolved’ patches required 255 PCs to account for 95% of variability.

### Machine Learning Classifier to Predict Mouse Model

Per mouse proportions of ‘Uninvolved’ patches and the ‘Involved’ k-means patch classes were generated for every sample from the archived mouse cohort and utilized to train a linear determinant analysis (LDA) classifier to predict the mouse model (Fig S5A). For inference, the final schematic is shown in Fig S5B. Overlapping patches are extracted from our prospective mouse cohort samples and filtered according to tissue maps generated by the small patch classifier. Our ‘Involved’ versus ‘Uninvolved’ classifier is applied, and ‘Involved’ patches undergo a further round of PCA-based dimensionality reduction that is first fit to archived mouse cohort data. Our k-means model, which is also trained on our archived mouse cohort, then categorizes ‘Involved’ patches into one of the discovered classes. Finally, the per mouse proportions of ‘Uninvolved’ patches and ‘Involved’ k-means patch classes are generated for each mouse and fed into the LDA classifier to infer the mouse model.

## Acknowledgements

The authors wish to acknowledge the Stony Brook Research Histology core for their expert assistance with paraffin embedding, sectioning, and performing H&E staining on our colon samples. The authors also wish to acknowledge the Stony Brook Division of Laboratory Animal Resources for their expert assistance with caring, housing, and maintaining mice used to generate results included in this manuscript. This work is support by grants from the National Institutes of Health: DK052230 to V.W.Y. and CA205109/CA225021 to J.H.S.

## Supporting Information Captions

**Supplemental Table 1.** Mouse Cohort Sample Numbers Per Genotype and Treatment.

|  | <u>Genotype</u> | <u>Injection</u> | <u># Days</u><br><u>Injection</u> | <u>DSS/H<sub>2</sub>O</u> | <u># Days</u><br><u>DSS/H<sub>2</sub>O</u> | <u>Abbreviation</u> | <u>Induction Status</u> | <u>Number</u> |
| --- | --- | --- | --- | --- | --- | --- | --- | --- |
| Archived Mouse Cohort | *Klf5*<sup>ΔIND</sup> | CO | 5 | N/A | 0 | 5C-*Klf5*<sup>ΔIND</sup> | Control | 2 |
|  | *Klf5*<sup>ΔIND</sup> | CO | 5 | H<sub>2</sub>O | 7 | 5C-*Klf5*<sup>ΔIND</sup> + H<sub>2</sub>O | Control | 9 |
|  | *Klf5*<sup>ΔIND/+</sup> | CO | 5 | H<sub>2</sub>O | 7 | 5C-*Klf5*<sup>ΔIND/+</sup> + H<sub>2</sub>O | Control | 3 |
|  | *Klf5*<sup>WT</sup> | CO | 5 | N/A | 0 | 5C-*Klf5*<sup>WT</sup> | Control | 2 |
|  | *Klf5*<sup>WT</sup> | CO | 5 | H<sub>2</sub>O | 7 | 5C-*Klf5*<sup>WT</sup> + H<sub>2</sub>O | Control | 6 |
|  | *Klf5*<sup>WT</sup> | TAM | 5 | N/A | 0 | 5T-*Klf5*<sup>WT</sup> | Control | 4 |
|  | *Klf5*<sup>WT</sup> | TAM | 5 | H<sub>2</sub>O | 7 | 5T-*Klf5*<sup>WT</sup> + H<sub>2</sub>O | Control | 5 |
|  | *Klf5*<sup>ΔIND</sup> | TAM | 5 | N/A | 0 | 5T-*Klf5*<sup>ΔIND</sup> | Colitis | 9 |
|  | *Klf5*<sup>ΔIND/+</sup> | CO | 5 | DSS | 7 | 5C-*Klf5*<sup>ΔIND/+</sup> + DSS | Colitis | 5 |
|  | *Klf5*<sup>ΔIND/+</sup> | TAM | 5 | DSS | 7 | 5T-*Klf5*<sup>ΔIND/+</sup> + DSS | Colitis | 3 |
| Prospective Mouse Cohort | *Klf5*<sup>ΔIND</sup> | CO | 5 | N/A | 0 | 5C-*Klf5*<sup>ΔIND</sup> | Control | 4 |
|  | *Klf5*<sup>ΔIND/+</sup> | N/A | 0 | N/A | 0 | *Klf5*<sup>ΔIND/+</sup> | Control | 2 |
|  | *Klf5*<sup>WT</sup> | N/A | 0 | N/A | 0 | *Klf5*<sup>WT</sup> | Control | 2 |
|  | *Klf5*<sup>ΔIND</sup> | TAM | 5 | N/A | 0 | 5T-*Klf5*<sup>ΔIND</sup> | Colitis | 5 |
|  | *Klf5*<sup>ΔIND/+</sup> | N/A | 0 | DSS | 7 | *Klf5*<sup>ΔIND/+</sup> + DSS | Colitis | 8 |
|  | *Klf5*<sup>ΔIND/+</sup> | TAM | 5 | DSS | 7 | 5T-*Klf5*<sup>ΔIND/+</sup> + DSS | Colitis | 3 |

**Supplemental Figure 1.**
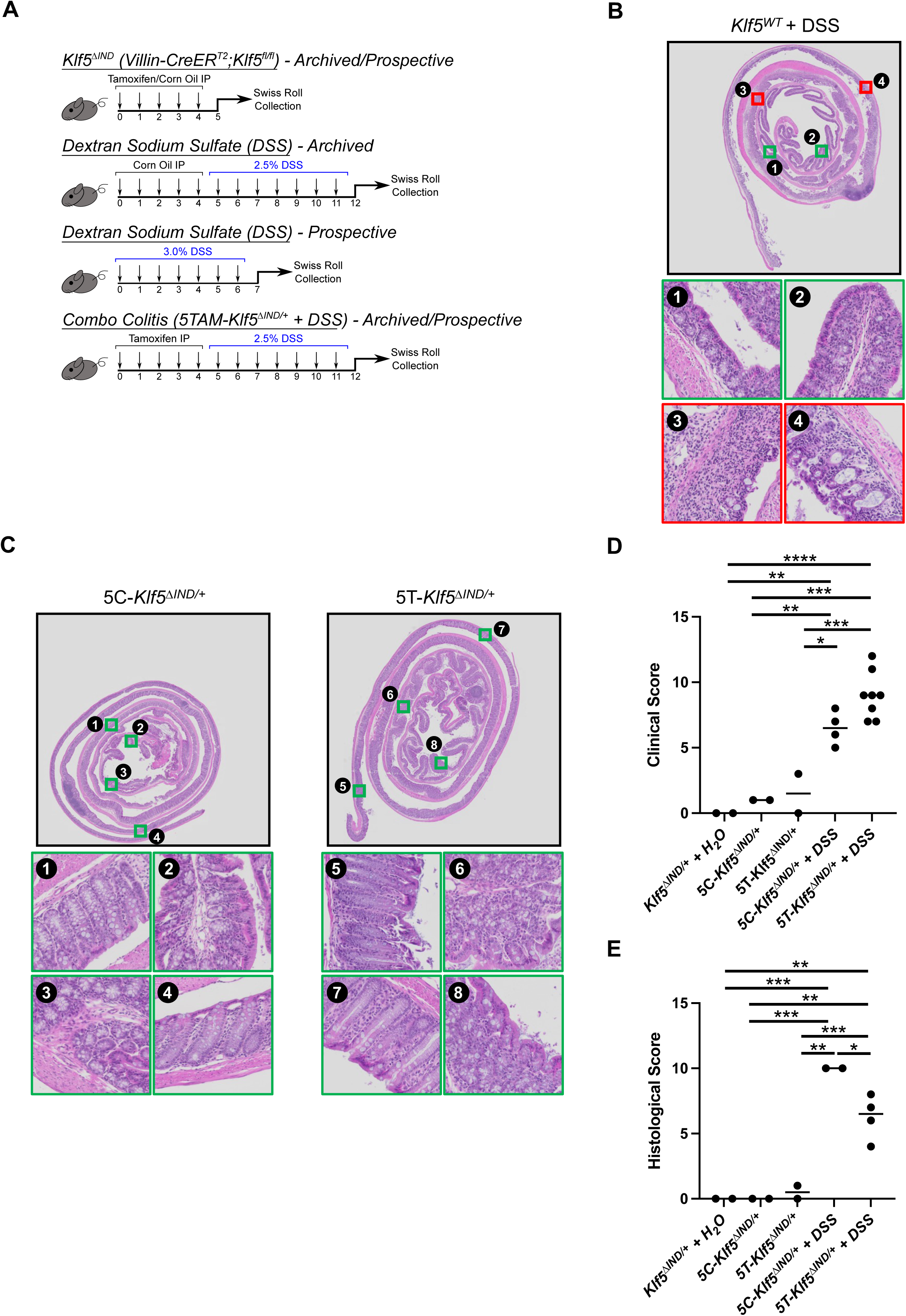
Colitis Mouse Models. A) Treatment schedules for each mouse model. DSS model in prospective cohort has no injections to confirm abnormal pathology is recognized independent of corn oil. Additionally, 3.0% DSS was used in prospective mice, as this was observed to be the optimal concentration at the Stony Brook facilities. B) Swiss roll of *Klf5*^*WT*^ (*Villin-CreER*^*T2*^*;Klf5*^*+/+*^) mouse treated with DSS and no TAM or CO injections. C) Though not utilized in this study, swiss rolls of corresponding controls (5T-*Klf5*^Δ*IND/+*^ and 5C-*Klf5*^Δ*IND/+*^) are shown for combined colitis model. D) Clinical scores combining weight loss, stool consistency, and fecal blood according to Cooper et al. (16) for combined colitis model, 5C-*Klf5*^Δ*IND/+*^ + DSS, and control mice with student’s t-test showing significant differences. E) Histological scores according to Cooper et al. (16). One-way ANOVA *p<0.05, **p<0.01, ***p<0.001

**Supplemental Figure 2.**
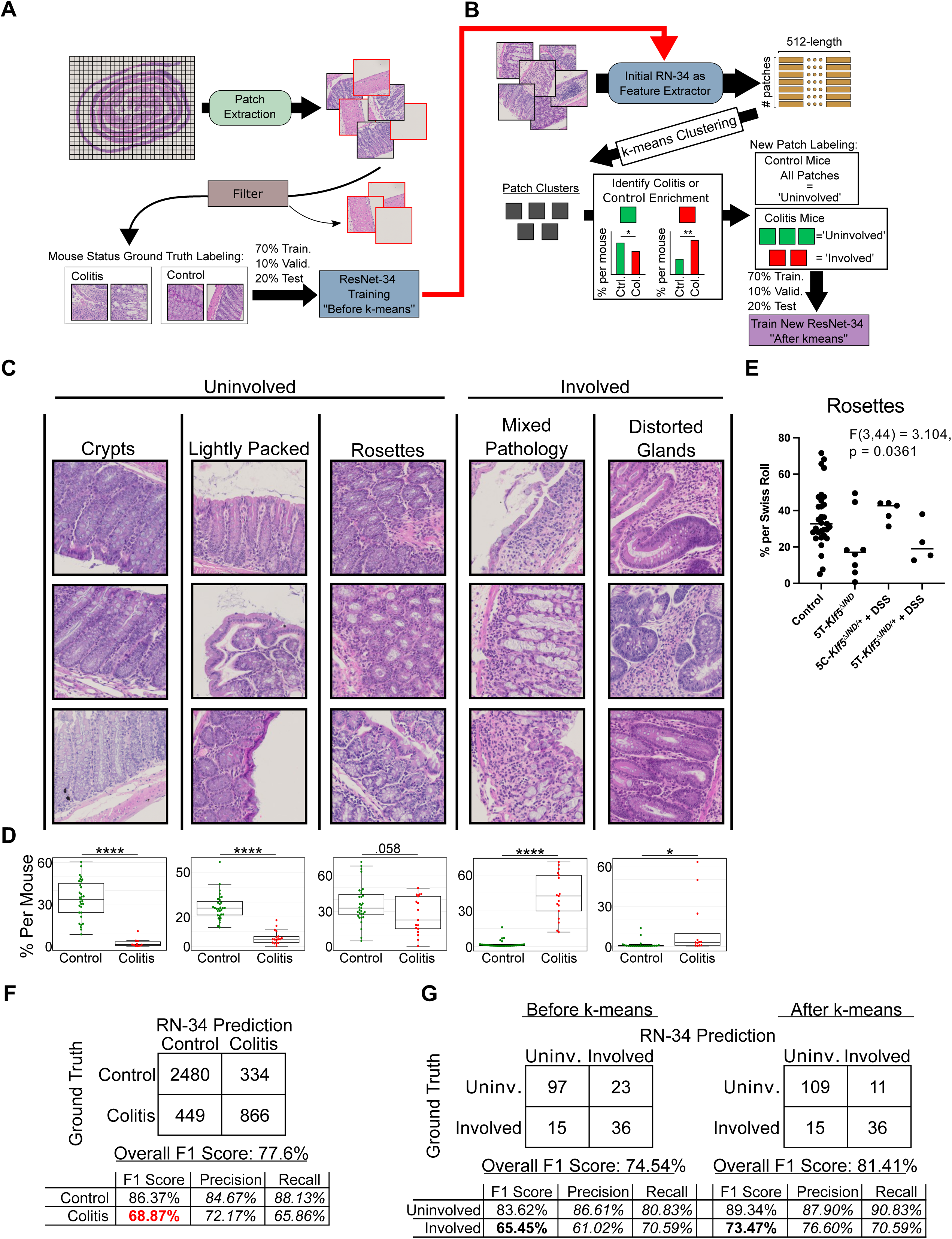
Second Phase of Training with ‘Involved’ and ‘Uninvolved’ K-Means Patch Cluster Labels Improves Model Performance. A) Overview schematic showing initial phase of RN-34 model training that uses mouse colitis status as patch ground truth labels. Thus, intracolonic heterogeneity in colitis mice is not addressed at this round of patch labeling. B) Second phase of H&E model training that uses trained RN-34 model from A) as a feature extractor for patches in dataset. K-means clustering is applied on extracted features to generate patch classes. Student t-test is utilized to assess whether patch classes are significantly enriched in proportion between colitis and control mice. In this second phase of training, all patches from control mice are labeled as ‘Uninvolved’, as no colitis induction was performed. For colitis mice, patches are given ‘Uninvolved’ or ‘Involved’ ground truth labels according to k-means predictions. C) K-means patch classes identified during second phase of training in B) used for ground truth labeling. D) Student’s t-tests show patch classes are found in differing portions between mice with colitis induction and controls. *p<0.05, **p<0.01, ***p<0.001, ****p<0.0001. E) Rosettes are in DSS-treated mice relative to other colitis models. One-way ANOVA shows a statistically significant difference between groups. F) Independent test set output confusion matrix for before k-means model. G) 200 patches from 4 mice (1 control, 1 of each colitis mouse model) were extracted and labeled as ‘Uninvolved’ or ‘Involved’ by a pathologist. 29/200 patches were discarded for not enough spatial context to provide a label. Inference using models trained in A) and B) show that the k-means patch labeling approach increased prediction agreement with pathologist-generated labels.

**Supplemental Figure 3.**
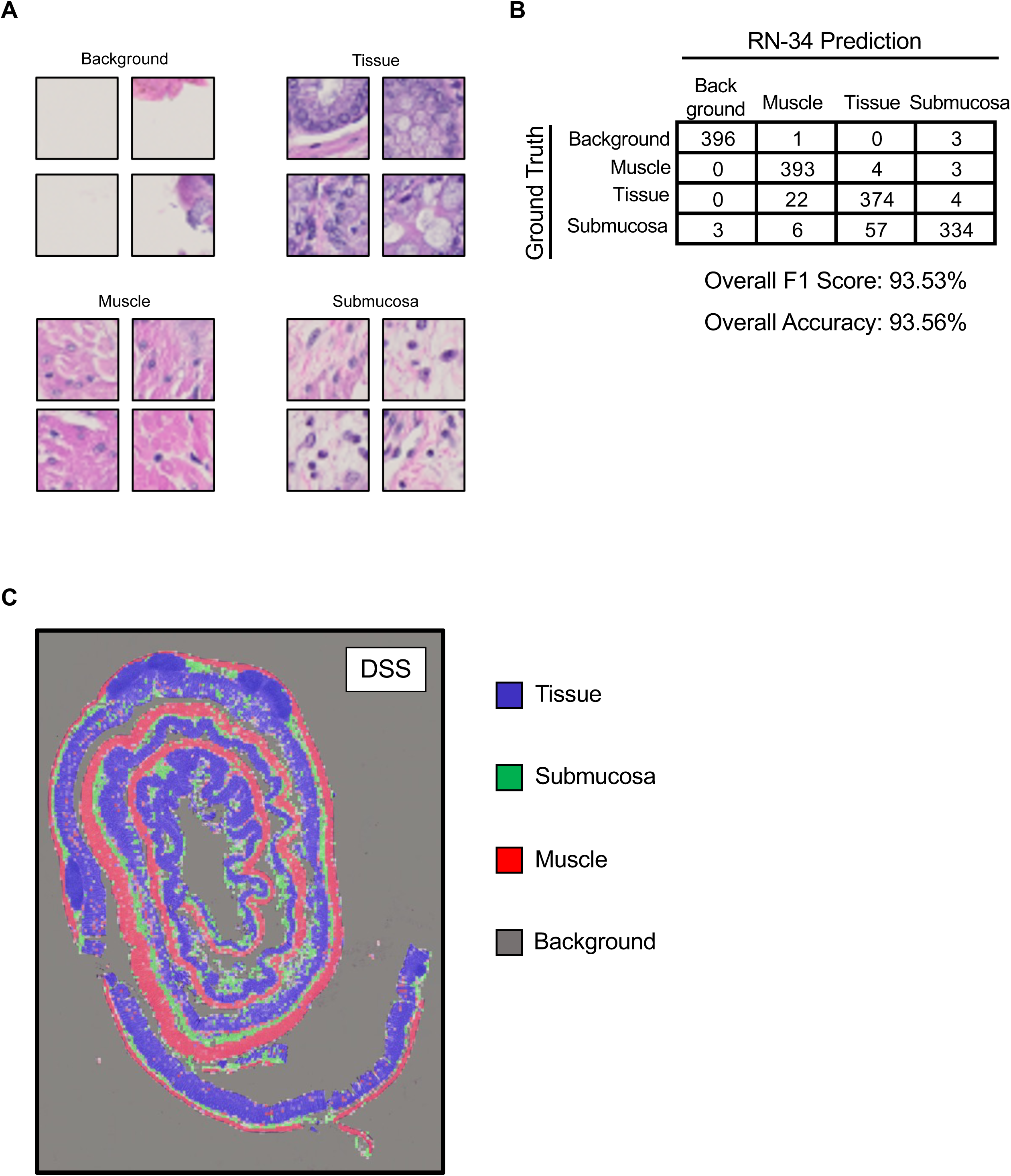
Small Patch Classifier Training for Preprocessing. A) Example 32×32 pixel patches for the ‘Background’, ‘Tissue’, ‘Muscle’, and ‘Submucosa’ classes used to train the small patch RN-34 classifier. B) Trained small patch RN-34 model confusion matrix outputs for independent test set of 4 mice, each with 100 patches of each class (1600 total patches). C) Example overlay of small patch RN-34 classifier on test set mouse.

**Supplemental Figure 4.**
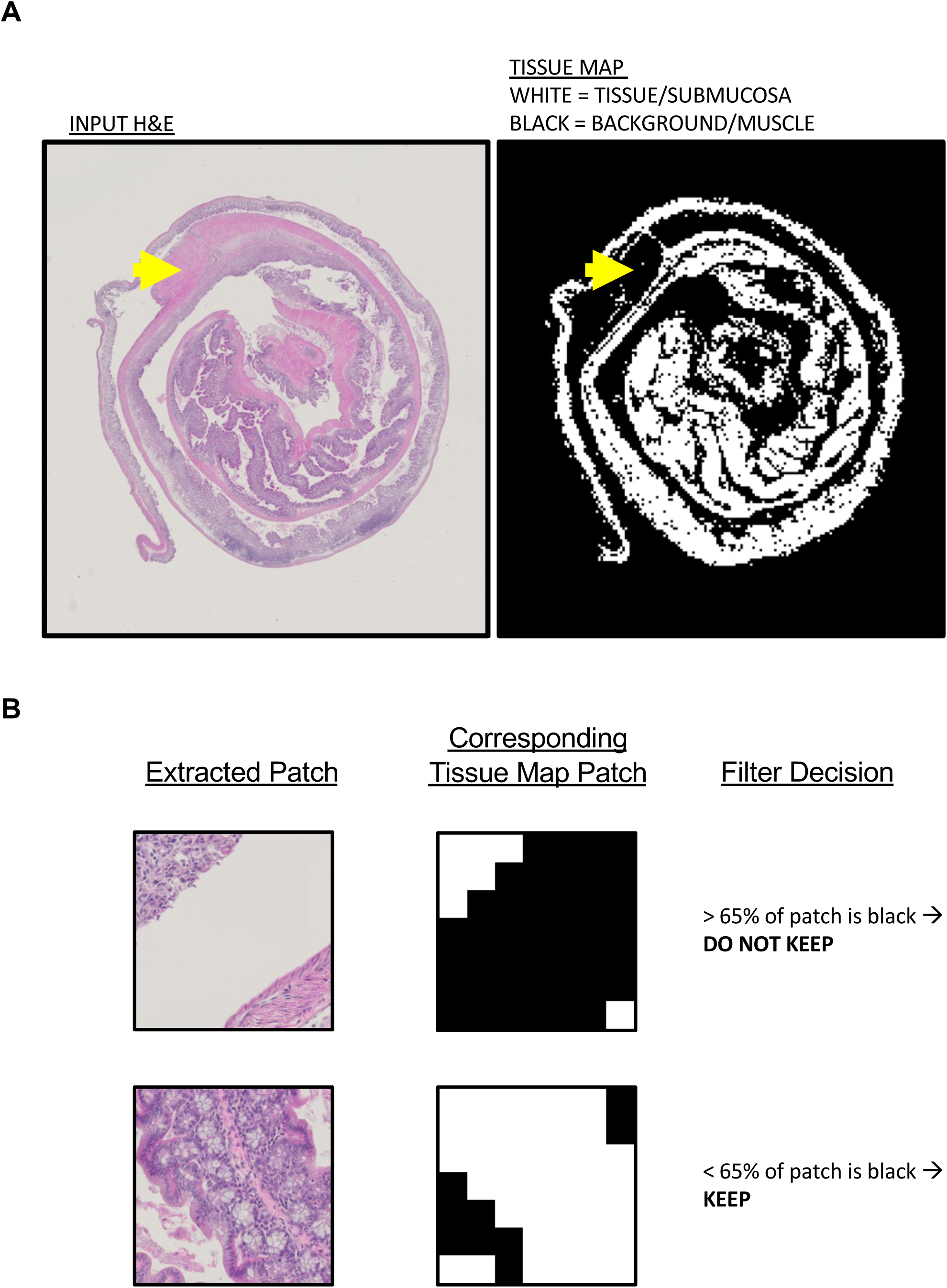
Tissue Map Generation and Filtering Process. A) Small patch RN-34 classifier from S4 Fig is applied to all 32×32 pixel patches extracted from a WSI. A tissue map is generated where ‘Background’ and ‘Muscle’ are black, while ‘Tissue’ and ‘Submucosa’ are white. The yellow arrow indicates a portion of muscle that is assigned to the Background/Muscle class on the corresponding tissue map. B) Example 224×224 pixel H&E patches with corresponding tissue map patches and filtering decisions. For each extracted patch, the decision to keep or filter out the patch is made based on whether there is more than 65% (filter) or less than 65% (keep) of unwanted Background/Muscle area on the corresponding tissue map patch.

**Supplemental Figure 5.**
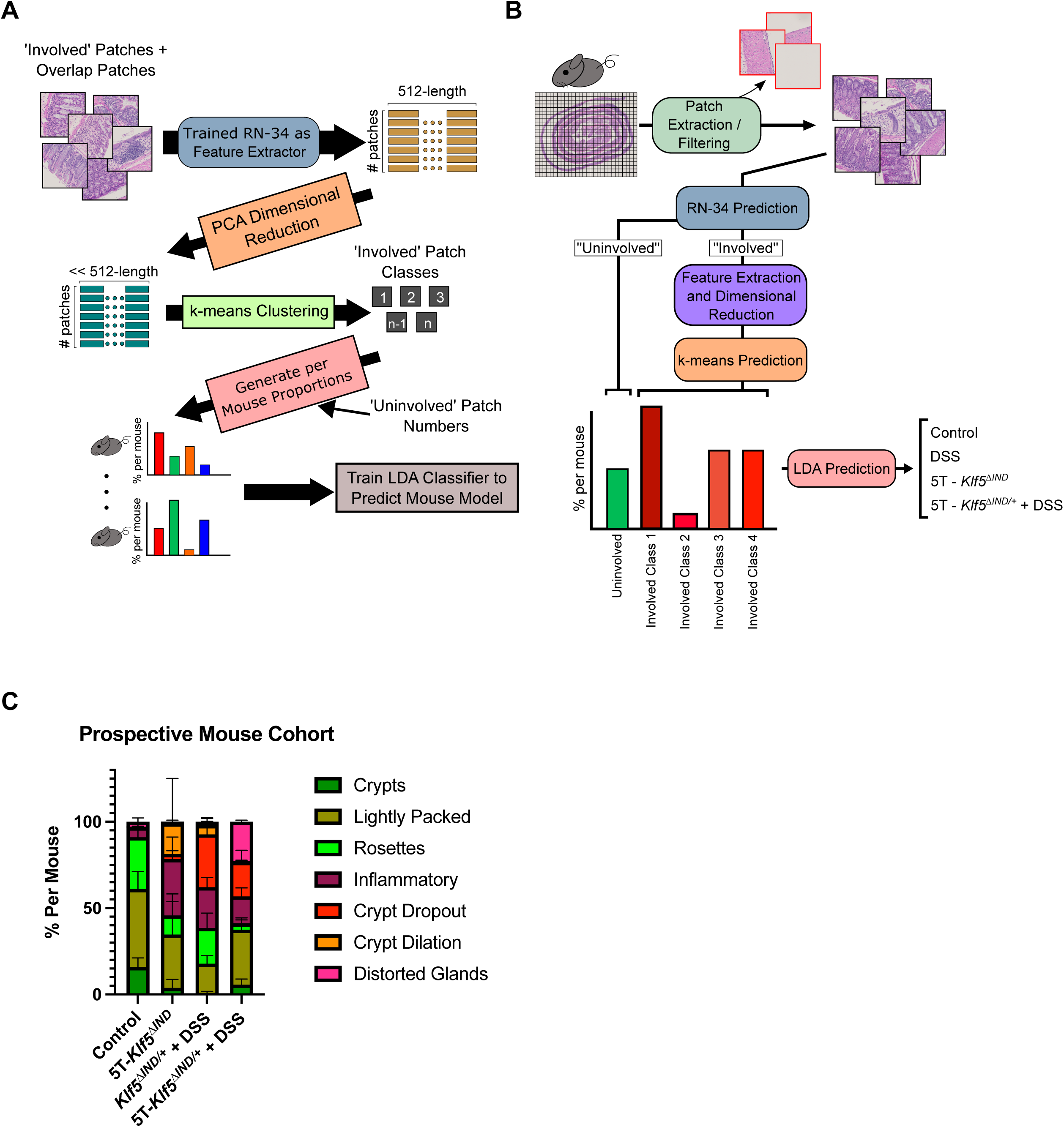
Linear Determinant Analysis Classifier Training and Inference Schematics. A) Overview schematic. All patches, including overlapping patches, classified as ‘Involved’ by our model undergo feature extraction by our trained ‘Involved’ versus ‘Uninvolved’ classifier. Subsequent PCA-based dimensionality reduction and k-means clustering identify 4 ‘Involved’ patch classes shown in Fig 4A. Per mice proportions of ‘Uninvolved’ patches and ‘Involved’ class patches are used to train an LDA classifier to predict mouse models. B) Overview schematic for inference pipeline. Overlapping patches are extracted and patches containing too much background or muscle are filtered out via tissue map process in S5 Fig. H&E classifier is applied to each patch to predict ‘Involved’ or ‘Uninvolved’. ‘Involved’ patches undergo further RN-34 feature extraction and PCA-based dimensionality reduction. K-means model trained on training cohort then classifies ‘Involved’ patches into one of the four ‘Involved’ patch classes. Patch proportion of ‘Uninvolved’ and ‘Involved’ k-means classes for each mouse are generated and fed into the LDA classifier for inference of mouse model. C) Stacked bar plot of total ‘Uninvolved’ and ‘Involved’ k-means patch class proportions across prospective mouse cohort. Error bars show mean with standard deviation.

**Supplemental Figure 6.**
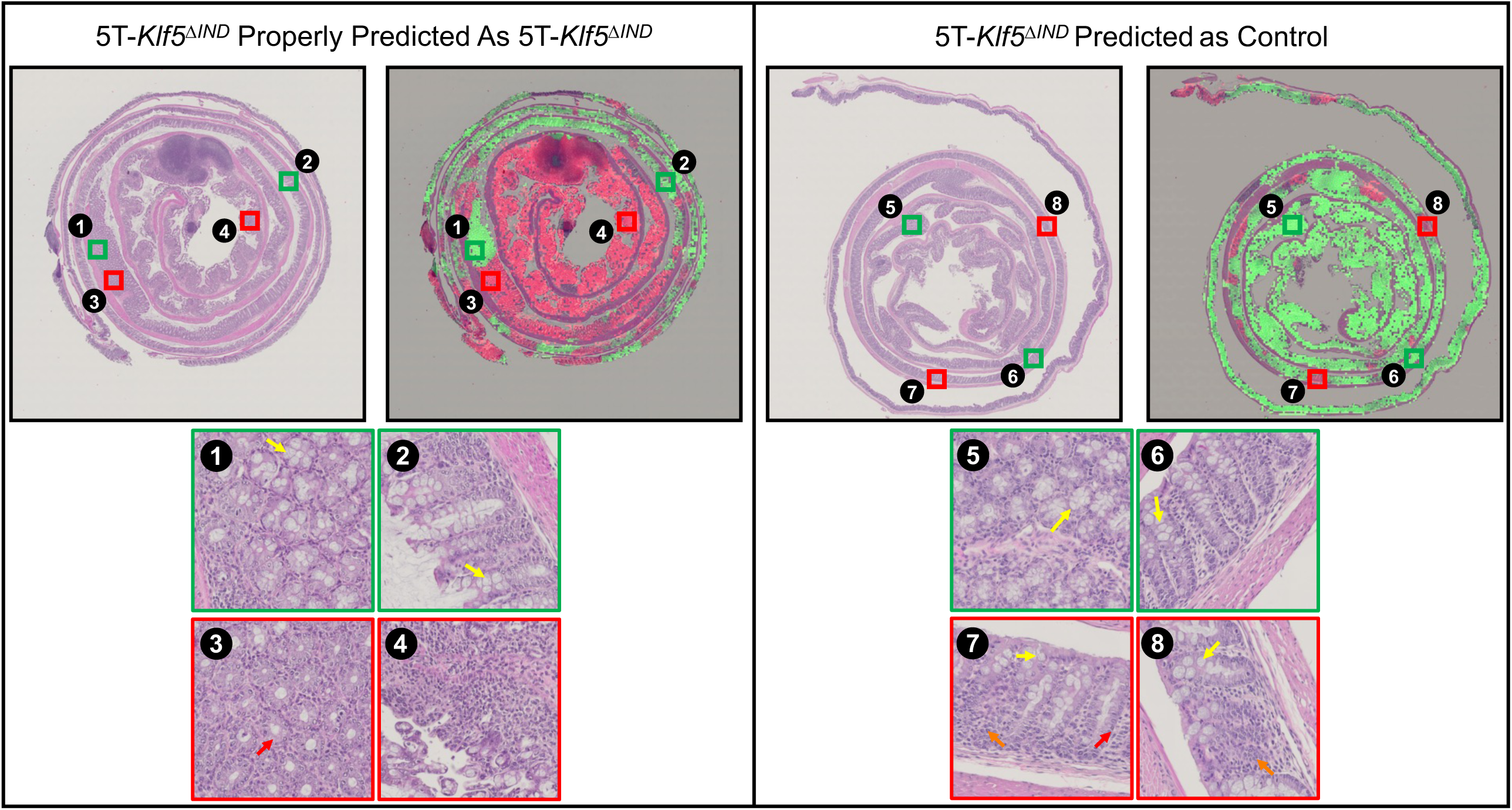
5T-*Klf5*^Δ*IND*^ Swiss Roll Predicted as Control has Healthier Appearing Histology Than 5T-*Klf5*^Δ*IND*^ Swiss Roll Predicted as 5T-*Klf5*^Δ*IND*^. Compared to the properly predicted 5T-*Klf5*^Δ*IND*^ swiss rolled colon (left), the colon predicted as control has fewer pathologies and likely represents a mouse with weak induction of colitis. Yellow arrows indicate presence of healthy goblet cells. Red arrows indicate loss of goblet cells. Orange arrows indicate crypt loss.

## Notes

### Competing Interest Statement

The authors have declared no competing interest.

### Summary of Updates

The supplemental table and figures have been added to the manuscript.

